# The influence of axial length upon the retinal ganglion cell layer of the human eye

**DOI:** 10.1101/2020.06.18.159871

**Authors:** Min Chen, Jill Nofziger, Ritobrato Datta, James C. Gee, Jessica Morgan, Geoffrey K Aguirre

## Abstract

**Purpose:** We examined the relationship between axial length and the thickness and volume of the ganglion cell layer (GCL) of the retina, and related these measures to the size of the optic chiasm.

**Methods:** We used optical coherence tomography to measure the thickness of the GCL over a 50° extent of the horizontal meridian in 50 normally-sighted participants with a wide range of axial lengths. Using a model eye informed by individual biometry, we converted GCL thickness to tissue volume per square degree. The volume of the optic chiasm was measured for 40 participants using magnetic resonance imaging.

**Results:** While GCL thickness decreases with increasing axial length, there is a positive relationship between GCL tissue volume and axial length, leading us to conclude that increasing axial length is associated with decreased retinal ganglion cell packing, increased cell size, or both. We characterize how retinal ganglion cell tissue varies systematically in volume and spatial distribution as a function of axial length. This model allows us to remove the effect of axial length from individual difference measures of GCL volume. We find that variation in GCL volume correlates well with the size of the optic chiasm as measured using magnetic resonance imaging.

**Conclusions:** Our results provide the volume of ganglion cell tissue in the retina, adjusted for the effects of axial length upon ganglion cell size and/or packing. The resulting volume measure accounts for individual differences in the size of the optic chiasm, supporting its use to characterize the post-retinal visual pathway.

## Introduction

The retinal ganglion cells receive signals that originate in the photoreceptors and transmit information to the central nervous system. These cells are located within the retinal ganglion cell layer (GCL) of the eye, along with displaced amacrine cells and supportive tissue. The technique of optical coherence tomography (OCT) has been regularly used to measure the thickness (in mm) of the GCL and other retinal layers. There is some interest in characterizing the GCL in healthy eyes, both to understand the contribution of these cells to normal vision and as a point of reference for measurements in ophthalmologic diseases that damage the retinal ganglion cells. A confounding property of OCT measurements in healthy control participants, however, is that the thickness of retinal tissue systematically varies with the size of the eye under study: the larger and more myopic the eye [1], the thinner the retina tends to be [2,3].

It may seem at first that this systematic variation of GCL thickness in eyes of different sizes should have an impact upon visual perceptual ability. The GCL is a relative bottleneck in the visual pathway, as the retinal ganglion cells are outnumbered over 100-to-1 by the both the photoreceptors and by the neurons of the primary visual cortex. Retinal ganglion cell density has been linked to the limit of grating acuity in the periphery of the visual field [4-8]. Specifically, the systematic decline in spatial acuity from parafoveal vision to the periphery follows very closely the corresponding decline in midget retinal ganglion cell receptive field density [6].

Given that the retina is thinner in larger eyes, we might therefore expect people with myopia, even with refractive correction, to have worse peripheral spatial acuity than emmetropes. This, however, is not the case. Measured with laser interference fringes, angular spatial acuity does not vary significantly as a function of individual differences in spherical refractive error [9, 10]. Because the limit of peripheral spatial acuity is plausibly linked to the density of retinal ganglion cells on the retina, these psychophysical findings suggest that eyes of different sizes nonetheless have on average the same number of retinal ganglion cells.

It is possible that the OCT and psychophysical measurements can be reconciled by considering the relationship between tissue thickness and volume in eyes of different sizes. The equivalence of spatial acuity across eyes of different sizes is only obtained when acuity is expressed in cycles per degree of visual angle (and when avoiding the effects of spectacle magnification from corrective lenses; [11]). In larger eyes, a greater surface area of retina (in square mm) subtends a given portion of the visual field (in square degrees of visual angle). It may be the case that eyes of different sizes have, on average, the same number of retinal ganglion cells, but that these cells are simply spread over a larger surface area in myopic eyes, resulting in a thinner retina. If we further assume that tissue thickness reflects only the number of cells present, with no changes in cell size or packing, then a prediction of this “stretching” model [11-14] is that measurements of the GCL in eyes of different sizes should be rendered equivalent when expressed as the volume of tissue that subtends a square degree of visual angle. An examination of this prediction, and a characterization of its failure, can contribute to efforts to derive estimates of retinal ganglion cell counts from OCT measures of the retinal layers [15-16].

Here we examine the relationship between axial length and the properties of the retinal ganglion cell layer of the human eye. We used OCT to measure the thickness of the GCL along the horizontal meridian in 50 normally-sighted participants. Biometric measurements from each participant were used to generate a model eye, which informed conversion of GCL thickness into tissue volume. Contrary to the prediction of the simple stretching model, we find that ganglion cell tissue volume is not constant across eyes of different sizes. This result suggests that cell packing and/or ganglion cell size vary systematically with axial length and contribute to total GCL volume. Using a principal component analysis, we explore the contribution that axial length has on the magnitude and spatial distribution of GCL tissue. The result is a model that captures individual differences in GCL tissue volume and thickness not attributable to variation in eye size. We considered that this measure may be more directly related to individual variation in the total number of retinal ganglion cells present in the eye. In support of this, we find that volumetric representations of GCL tissue best account for individual variation in the size of the optic chiasm, suggesting that this measure is appropriate for considering properties of post-retinal visual processing.

## Methods

### Ethics Statement

This study was approved by the University of Pennsylvania Institutional Review Board, and all participants provided written consent.

### Participants

We studied 50 normally sighted participants, recruited from the University of Pennsylvania and the surrounding Philadelphia community. Participants were required to be at least 18 years of age, have no history of ophthalmologic disease, and have corrected visual acuity of 20/40 or better. The median age of the participants was 26, and all but three participants were below the age of 40. There were multiple sessions of data collection for each participant, typically across multiple days. These sessions included several retinal imaging studies, as well as structural and functional magnetic resonance imaging. After measurement of distance acuity, participants underwent pharmacologic dilation of the pupil prior to collection of biometric measures and retinal imaging. Demographic and biometric information for the participants is provided in Table 1.

**Table 1.** Demographic and biometric data. We studied 50 subjects. The median and inter-quartile range are provided for several measures. SR = spherical refractive error obtained by auto refraction.

| Age<br>[years] | # Male /<br># Female | Acuity<br>[log MAR] | SR<br>[Diopters] | Axial length<br>[mm] | K1<br>[Diopters] | K2<br>[Diopters] |
| --- | --- | --- | --- | --- | --- | --- |
| 26 ± 10.1 | 26 / 24 | 0.05 ± 0.068 | -1.125 ± 4.25 | 24.1 ± 2.03 | 43.4 ± 2.32 | 44.0 ± 2.05 |

### Measurement of visual acuity

Best corrected distance visual acuity was measured for both eyes for every participant using the Early Treatment of Diabetic Retinopathy Study (ETDRS) chart [17]. The logMAR acuity for each eye was calculated, with interpolation between lines based upon number correct [18]. For 48/50 participants, acuity between the two eyes was within one line. We therefore report the average acuity of the two eyes.

### Biometry

All 50 participants underwent auto-refraction of both eyes using an auto kerato-refractometer (Topcon). We measured the axial length of both eyes in all 50 participants using the IOLMaster 500 (Zeiss). This instrument was also used to obtain keratometric measurements of both eyes in 40/50 participants. There was a high correlation of these measurements between eyes across participants (all Pearson’s correlation values > 0.95). Therefore, we took the average of the values across eyes in creating a model eye for each participant (after mirroring the measure of corneal torsion). For the ten participants who did not undergo keratometry, we assumed the mean measures obtained in the remainder of our population, which were 43.40 diopters at 0 degrees, and 44.34 diopters at 90 degrees.

### Optical coherence tomography

We imaged both eyes of every participant using the Heidelberg Spectralis OCT. The imaging session included several measurements; we describe here only the measurements made on the horizontal meridian. For each eye, three overlapping acquisitions of 30° width were made along the horizontal meridian. Each acquisition was a high resolution horizontal line scan that included the fovea. Retinal tracking was enabled on the device and the operator saved the scans after 100 individual b-scans were automatically aligned and averaged using the built-in Spectralis software. The images were exported from the instrument in DICOM format for further analysis, and retained their original x-axis position values in units of degrees of visual angle. The measurement of the left eye of one participant used high speed rather than high resolution acquisition settings. Therefore, only the high-resolution images of the right eye for this participant were used in the subsequent analyses.

### OCT montage

For each eye for each participant we created a montage of the three, overlapping horizontal line scans. Using the central fovea scan as a reference, we performed a global rigid registration between the other two scans onto the central scan. Our registration approach used a standard SIFT [19] feature detection on the intensity of each image, and a RANSAC [20] based optimization for aligning the features and images. Figure 1 shows the individual B-scans on the top row, and the montaged full scan on the bottom row.

**Figure 1.**
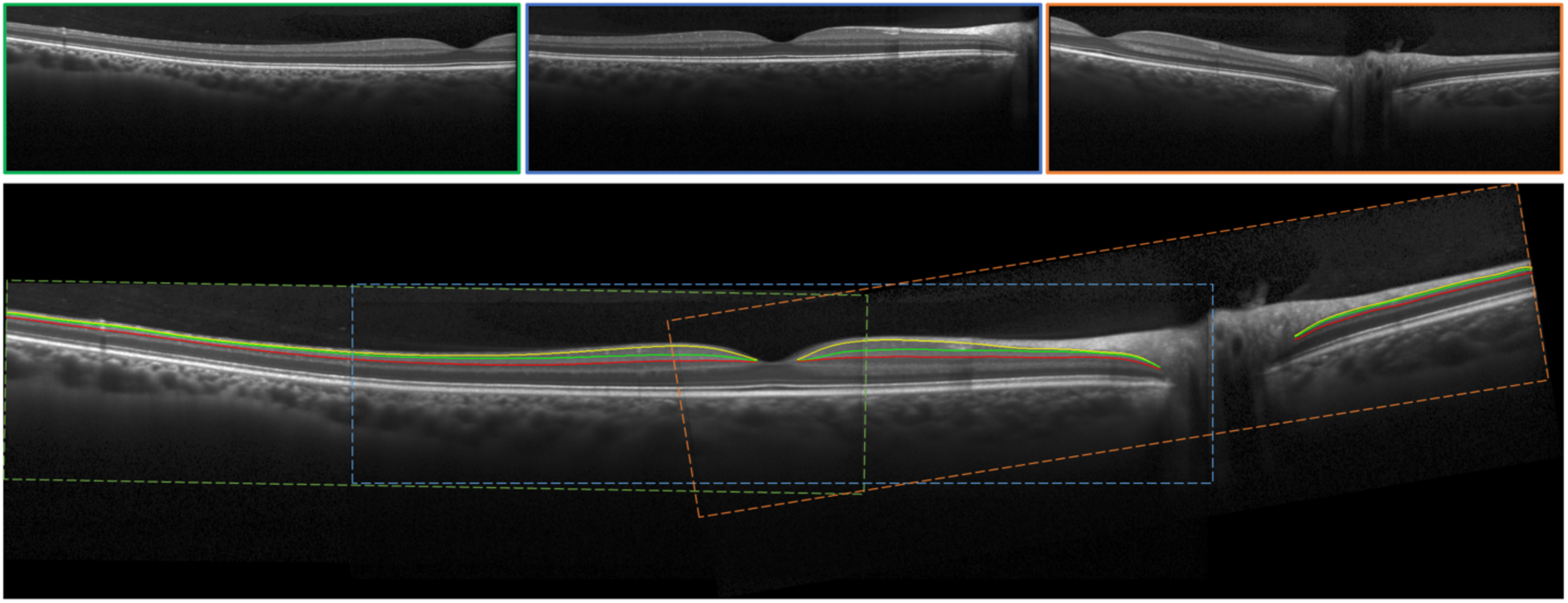
Example horizontal line scans. Three images (top row) each covering 30 degrees of retina were acquired for each eye for each subject and then montaged together into a single image (bottom row). The GC and IP layers were manually labeled. The borders (yellow, green, red) defined by these labels were smoothed with a spline.

### GC layer segmentation and measurement

We performed manual segmentation to identify the ganglion cell (GC) and inner plexiform (IP) layers. The montaged horizontal line images were viewed using ITKSNAP [21]. One of the authors (JN) labeled pixels on the image as corresponding to the GCL or IPL. A third category (“indistinct”) was assigned to regions of the GC and IP layers where the boundary between the two could not be discerned, typically as the result of “shadow” artifacts from overlying retinal blood vessels. The 3 aligned images in each montage were observed simultaneously, and the best defined layer in the three images was used for the segmentation.

The manual segmentation implicitly defined three boundaries: 1) The retinal nerve fiber layer with the GCL; 2) the GCL with the IPL; and 3) the IPL with the inner nuclear layer. We fit 15^th^ order cubic splines to these borders, allowing us to interpolate across short patches in which the boundary between the GC and IP layers was indistinct. As each boundary is discontinuous at the fovea and optic nerve, three disjoint splines were individually fit to the following three sections of the retina: 1) from the temporal extent of the image to the fovea center, 2) from the fovea center to the optic nerve, and 3) from the optic nerve to the nasal extent of the image.

Using the smoothed delineations, we obtained the thickness of the GC and IP layers at equal eccentricity positions along the GC/IP boundary. At each location, we calculated the closest Euclidean distance from the GC/IP boundary to the inner GC and outer IP boundaries, thus providing the thickness (in mm) of each layer.

The layer measurements showed a high degree of similarity between left and right eyes for each participant. We found that the Pearson’s correlation of the mean thickness (across eccentricity) of the GC+ IP layers, between left and right eyes across participants, was R = 0.89. This high correlation indicates that measurements from each eye are not independent, and suggests that combining the measurements from the two eyes within participant will yield a better estimate of individual participant variation. Therefore, in subsequent analyses, we averaged the measurements between the two eyes for each participant. To do so, we mirror-reversed the GC and IP thickness profiles from the left eye, aligned the measurements between the two eyes at the fovea, and averaged the values.

### Calculation of retinal area per square degree

A software model eye [22] was used to calculate the relationship between the retinal surface and visual field. The model was customized for each participant using their biometric information, including axial length, spherical refractive error, and keratometry values (when available). Variation in the overall size and dimensions of the vitreous chamber was derived largely from David Atchison’s work [23]. Ray tracing through the model provided an estimate of mm^2^/degree^2^ at positions along the horizontal meridian, passing through the fovea. These values were then used to convert the thickness profiles into the volume of tissue per square degree of visual angle.

### PCA approach

We applied a principal component analysis (PCA) to the profiles of GCL tissue volume across participants. Due to individual differences in the extent of the fovea and position of the optic disc, some eccentricity points had measurements from a subset of the 50 participants studied. We confined the PCA analysis to those eccentricity locations in which 45/50 participants had a thickness measurement. The PCA was conducted in MATLAB using the alternating least squares algorithm to impute the values at those locations with measurements from more than 45 but fewer than 50 participants. The resulting PCA scores were observed to have small areas of high variation that were suspected to reflect noise. To reduce the influence of these, we conditioned the scores with a cubic smoothing spline (smoothing parameter = 0.1). All subsequent calculations of variance explained and fits to the data were conducted with these smoothed scores.

### Measurement of optic chiasm volume

We collected structural brain MRI images from 40 of the participants. MRI scanning made use of the Human Connectome Project LifeSpan protocol (VD13D) implemented on a 3-Tesla Siemens Prisma with a 64-channel Siemens head coil. A T1-weighted, 3D, magnetization-prepared rapid gradient-echo (MPRAGE) image was acquired for each participant in axial orientation with 0.8 mm isotropic voxels, repetition time (TR) = 2.4 s, echo time (TE) = 2.22 ms, inversion time (TI) = 1000 ms, field of view (FoV) = 256 mm, flip angle = 8°.

The MRI data were processed using the Human Connectome Project minimal processing pipeline [24]. This pipeline includes application of the FreeSurfer v5.3 toolbox, which performs tissue segmentation of the anatomical data [25-28]. The optic chiasm was automatically identified and demarcated in the MPRAGE image for each participant in their native anatomical space based on registration to a probabilistic atlas in FreeSurfer [29].

### Data and code availability

The code used to analyze the data and produce the figures is publicly available (https://github.com/gkaguirrelab/retinaTOMEAnalysis). The data presented in the paper will be made available for download following publication.

## Results

We obtained OCT and eye biometric measurements from 50 normally-sighted participants. Table 1 provides the demographic and biometric information for this group. Spherical refractive error varied substantially across the group, ranging from –10.25 D to +3.88 D. Axial length also varied substantially, from 21.8 to 27.6 mm. Consistent with prior reports [2], there was a strong negative correlation between axial length and spherical refractive error across our participants (R=–0.83). Despite this marked variation in refractive error, all participants had corrected, central distance acuity of –0.0678 logMAR (decimal equivalent 20/23) or better. There was no relationship between axial length and best-corrected foveal, distance visual acuity (Pearson’s R = 0.07).

### Relationship between GCL thickness, volume and axial length

We collected horizontal OCT line scans through the fovea from both eyes of our participants (Figure 1, top). These acquisitions were montaged, segmented, and combined across eyes to obtain the thickness profile of the retinal ganglion cell layer (GCL) for each participant across a 50° extent (Figure 1, bottom).

Figure 2a presents the GCL thickness profiles for each of the 50 participants, along with the group mean. The form of the thickness profile is in good agreement with prior reports [30,31], including the finding of a thicker parafoveal region in the nasal as compared to temporal retina. There is substantial individual variation in the profiles, most notably in the peak thickness of the temporal parafoveal region, which ranged from 0.04 to 0.06 mm. We obtained the mean thickness of the GCL profile across eccentricity for each participant, and compared this thickness value to axial length (Figure 2b). We find a modest negative correlation (R=–0.24), consistent with similar measurements of the inner retina [3, 32-35].

**Figure 2.**
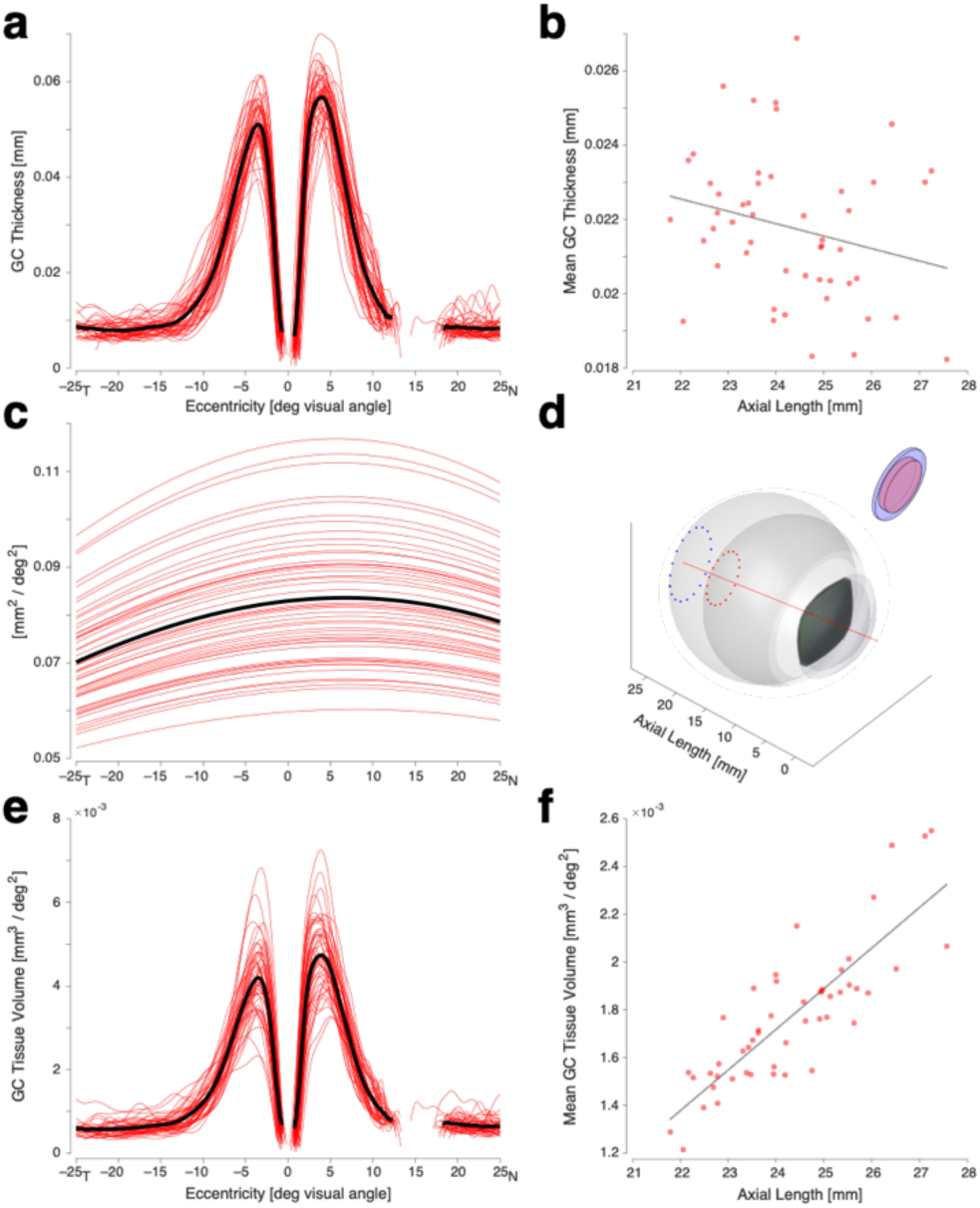
Relationship between GC layer thickness, volume, and axial length. (a) GC thickness profiles for 50 subjects (red) and the population mean (black), extending from –25 degrees (temporal) to 25 degrees (nasal). (b) The relationship between axial length and the mean GCL thickness from each participant. (c) The calculated retinal surface area (in mm2) that subtends a square degree of visual angle as a function of horizontal position on the retina. The function varies across subjects (red) primarily due to differences in axial length. The population mean is shown in black. (d) Rendering of the model eyes generated for the participants with the shortest and longest axial length. The dotted circles subtend a 30° diameter of the visual field in each eye around the visual axis (red line). Illustrated inset is a patch of retina from these two eyes that each have a diameter of 30°, have the same tissue volume, and correspondingly have different thickness. (e) The volume of GC tissue per square degree of visual angle as a function of retinal eccentricity for the 50 participants (red) and the population mean (black). (f) The relationship between axial length and the mean GC tissue volume per square degree across eccentricity from each participant.

One source of variation in retinal thickness may simply be differences in retinal surface area in eyes of different sizes. This possibility has been described as the “stretching” account of retinal development (e.g., [11-14]). We used a software model eye [22], informed by individual biometric measurements, to simulate vitreous chamber size and the optics of the eye for each participant. The simulation provides the retinal area (in mm^2^) that subtends the visual field (in degrees^2^) as a function of position along the horizontal meridian of the eye (Figure 2c). The form of this conversion across eccentricity for the emmetropic eye is in good agreement with the equations of Drasdo and Fowler (1974) [36], and as updated by Watson (2014) [37]. As expected, substantial variation in the conversion factor is present across participants, attributable to variation in axial length (Figure 2d).

We applied the conversion functions to the GCL thickness profile from each participant. The result is a profile of the volume of GCL tissue (in mm^3^) that subtends a square degree of visual angle (Figure 2e), expressed as a function of eccentricity in visual degrees. The general form of the profile is unaltered by this conversion, although we observe that the variation in the overall tissue volume across participants is greater than the variation that was observed in layer thickness. If it is the case that there is no systematic relationship between axial length and the total number of retinal ganglion cells present in the eye, then we might expect this volumetric representation of the data to no longer be related to axial length. However, when we examine the relationship between the mean (across eccentricity) tissue volume and axial length, we find that the conversion has introduced a strong, positive correlation between axial length and tissue volume (R=0.84) (Figure 2f).

This result indicates that variation in axial length has influences upon the GCL beyond simply the area with which the equivalent number of cells is spread over the retinal surface. Specifically, larger eyes have larger retinal ganglion cells, lower cell packing density, or both. Given this, if we wish to relate OCT measures of layer thickness to retinal ganglion cell number independent of incidental factors (cell size, packing), then it is necessary to characterize how axial length influences the overall volume and distribution of GCL tissue.

### Characterization model for GCL volume profiles

Beyond establishing a relationship between axial length and mean GC tissue volume, we wished to characterize how variation in axial length influences the profile of GC tissue across the horizontal meridian. If we are able to model the effect of axial length upon the GCL profile, then we can correct the GC profile for the effect of axial length.

To do so, we first conducted a principal components analysis upon the tissue volume profiles of our 50 participants (Figure 3). To reduce the influence of noisy measurements from eccentricity locations with missing values from some participants, we conditioned the principal components with a cubic smoothing spline. We found that the first 6 components were sufficient to explain 99.5% of the variance in the data. The first component primarily represents the mean tissue volume profile. Subsequent components capture variation in the distribution of GC tissue across eccentricity. For example, components 2 and 4 characterize the relative tendency of tissue to be crowded towards the parafoveal hills or pushed more towards the periphery. Figure 4 shows the reconstruction of individual participant tissue volume profiles using only the first six, smoothed, principle components. We find that this low-dimension representation well represents the original profiles of GC tissue volume.

**Figure 3.**
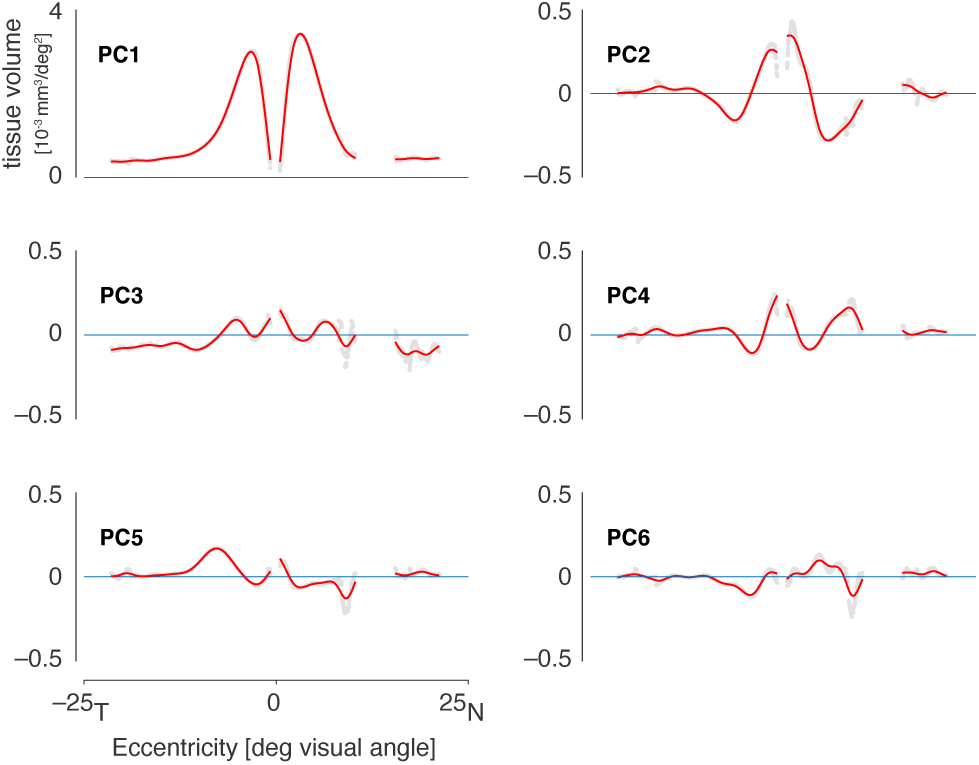
The first six principal components derived from an analysis of GC tissue volume profiles across the 50 participants. The initial (gray) components were smoothed (red) to reduce the influence of noisy regions of the measurement.

**Figure 4.**
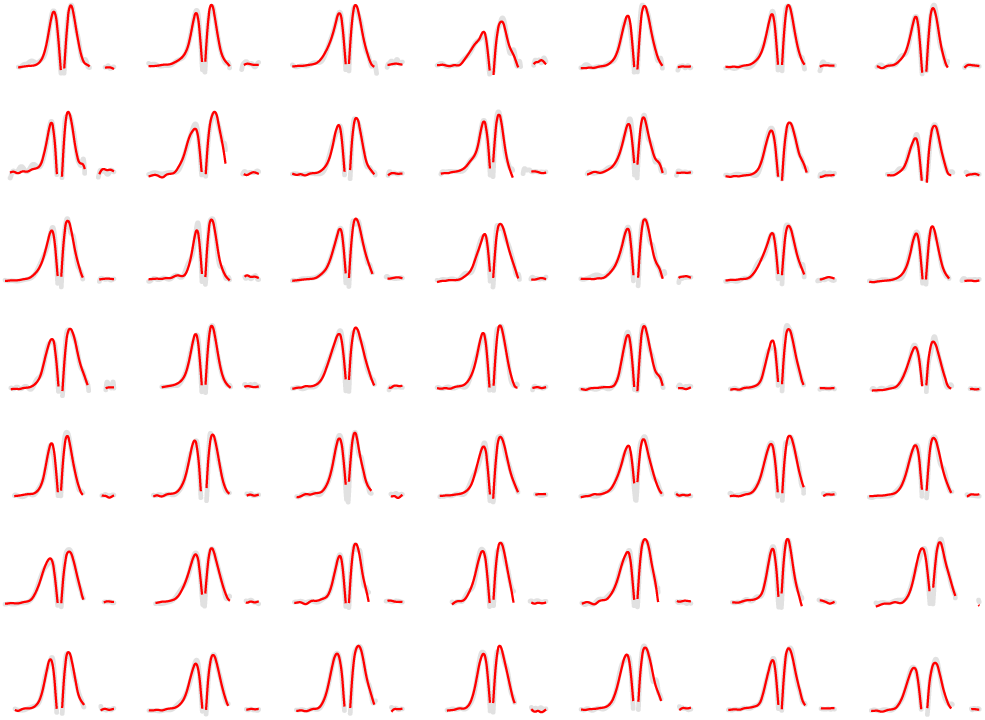
Original (gray) and reconstructed (red) GC tissue volume profiles for the first 49 participants, using the first 6 smoothed principle components, demonstrating the accuracy of the low dimensional representation of the data.

The GC tissue volume profile for each participant can be described by the loading on the six components. We examined if variation in axial length produces systematic variation in these weights. We performed a regression of each principal component loading with respect to axial length, which creates a hyperplane whose normal vector describes the direction and magnitude that axial length has on each principal component. We find a significant and substantial relationship with axial length for PC1 and PC2, and a weak, albeit significant, relationship with PC4. Table 2 summarizes the properties of the principal components.

**Table 2.** Properties of the principal components derived from the GCL tissue volume profiles across subjects. The first six principal components were found to explain 99.5% of the total variance in the data. The first principal component primarily accounts for variation across subjects in the mean tissue volume across the profile. Components 2-6 account for 75% of the remaining variation in the shape of the profile across subjects. The Pearson correlation of axial length with the weights on each of the components is also given.

| PC | Shape variance explained | Correlation w/ axial length |
| --- | --- | --- |
| 1 | -- | 0.85 |
| 2 | 40% | 0.75 |
| 3 | 12% | 0.11 |
| 4 | 10% | 0.25 |
| 5 | 8% | -0.16 |
| 6 | 5% | -0.13 |

These results may be used to visualize the effect of axial length upon the GCL. Figure 5 presents (in green) the GCL profile for an emmetropic eye, and then how that profile is altered with variation in axial length for each of the six principal components. As can be seen, variation in axial length scales the overall tissue volume (PC1), but also adjusts the distribution of tissue across eccentricity (PC2, and minimally PC3 and 4). In myopic eyes, retinal ganglion cell tissue is crowded more closely towards the fovea.

**Figure 5.**
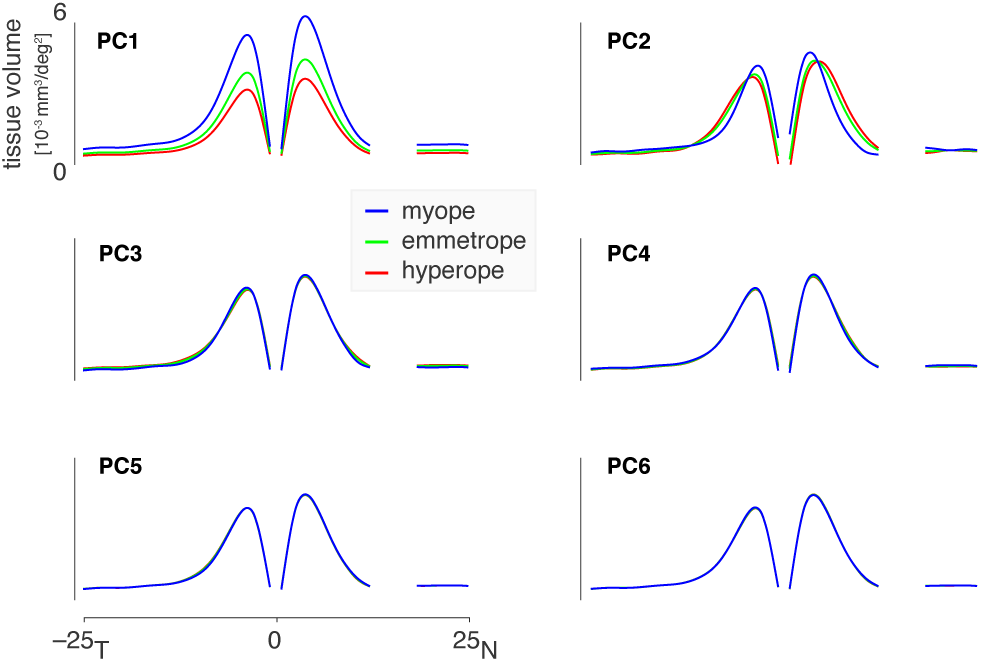
The influence of axial length on the GC tissue volume profile, as reflected in each of the first 6 principal components. Each plot shows how one of the principal components varies with axial length from myopic (blue, 27.57mm), to emmetropic (green, 23.58mm), to hyperopic (red, 21.79mm).

Having characterized the effect of axial length upon the GCL tissue volume profile, it is now possible to adjust the profile obtained for any one participant to remove the influence of axial length. In effect, we can express the GCL tissue volume profile for each participant, projected to the emmetropic eye. Figure 6a presents the individual and mean profile following this adjustment. While individual differences remain, the extreme variation seen in Figure 2e has been removed. Figure 6b confirms that the effect of axial length in the data has been removed.

**Figure 6.**
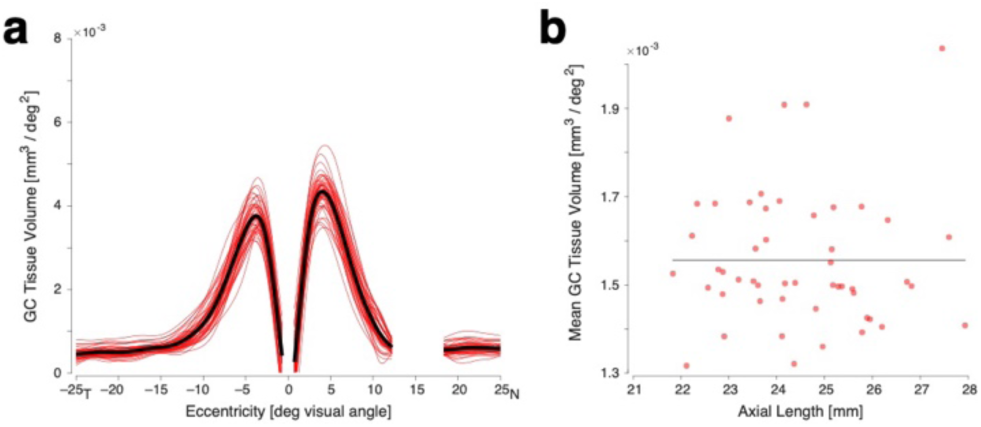
(a) GC tissue volume profiles for the 50 subjects after removing the effect of axial length by projecting each profile to the emmetropic eye. (b) The relationship between axial length and the mean GCL tissue volume per square degree across eccentricity from each participant following correction for variation in axial length.

Individual differences in the overall volume of retinal ganglion cell layer tissue remain after adjustment for the effect of axial length. We next asked if this adjusted measurement may be related to other properties of the visual pathway that reflect the influence of retinal ganglion cells.

### Relation to optic chiasm volume

The retinal ganglion cells give rise to axons that form the optic nerve and decussate at the optic chiasm. We considered that individual differences in the size of the optic chiasm would reflect individual differences in the number of retinal ganglion cells in the retina. We obtained a high resolution, T1-weighted image of the brain in 40 of the 50 participants, and derived from this image the volume of the optic chiasm. We then examined the correlation across s between optic chiasm volume and retinal measures.

We first examined the relationship between mean GCL thickness (Fig 2b) and optic chiasm volume, and found a significant relationship (Pearson’s R = 0.37 ± 0.20 SEM obtained by bootstrap resampling across the participants, p = 0.0173). Next, we examined this relationship for mean GC tissue volume, unadjusted for axial length (Fig 2f). This correlation was substantially higher (Pearson’s R = 0.51 ± 0.11 SEM, p = 6.8e-4), indicating that the volume of ganglion cell tissue is a better predictor of the size of the optic chiasm than is the thickness of this retinal layer. We then examined individual differences in the volume of ganglion cell tissue, after adjusting for the modeled effects of axial length (Fig 6b), and found that this measure is also strongly related to optic chiasm size (Pearson’s R = 0.53 ± 0.16 SEM, p = 4.5e-4).

The adjusted and unadjusted volume measures are correlated with one another (Pearson’s R = 0.50), but presumably differently reflect properties of the ganglion cell layer of the eye. By appeal to results from psychophysics, we propose that the GCL volume measure after adjustment for axial length reflects the number of retinal ganglion cells. In contrast, the unadjusted volume is dominated by variation in cell size and packing. We find that both volume measures (adjusted and unadjusted) are well correlated with the size of the optic chiasm. We therefore examined if the adjusted and unadjusted volume measures each make an independent contribution to the prediction of optic chiasm size. We included both measures in a linear model of optic chiasm size. Overall, this model explained significant variance in the optic chiasm measurements [F(2,40) = 10.9, p = 0.00019]. Further, both the adjusted and unadjusted GCL volume measures made significant, independent contributions to the model (p = 0.022 and p = 0.029, respectively). The relative weighting of the two GCL measures was effectively equal. Figure 7 presents the performance of this combined model. On the x-axis is the combined score for each participant of the two GCL volume measures (expressed as the deviation of each subject from the across-participant mean of this score), and on the y-axis the optic chiasm volume. Those subjects who tended to have a greater volume of GCL tissue, tended to have a larger optic chiasm.

**Figure 7.**
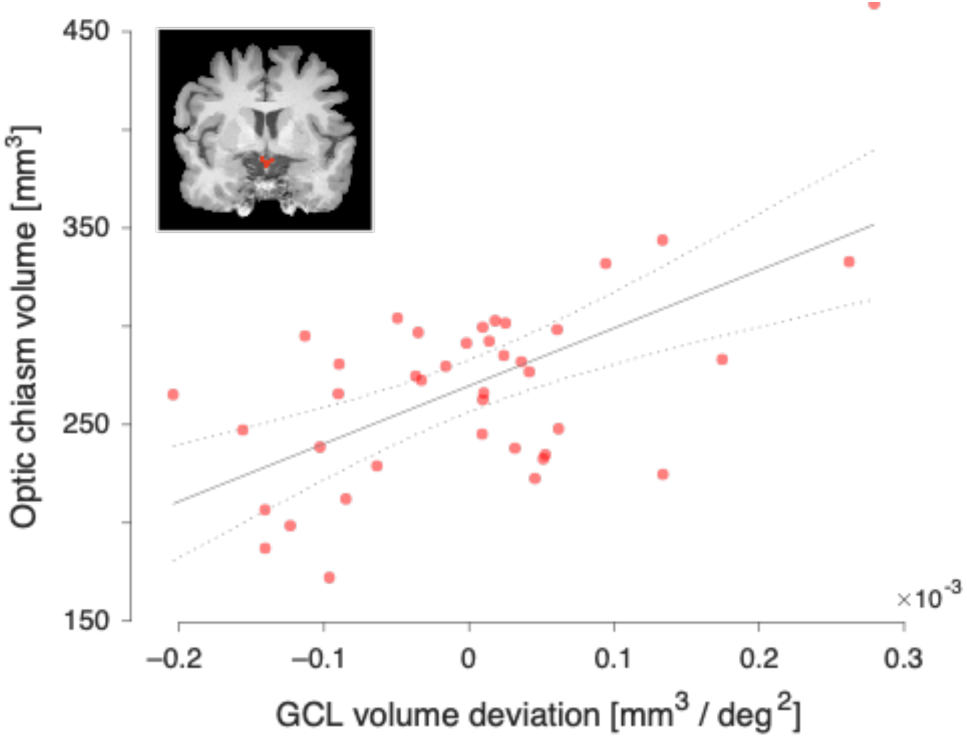
Relation of modeled GC tissue volume to optic chiasm volume. Optic chiasm volume was measured from an anatomical MRI image in each of forty participants (example coronal MRI image inset; optic chiasm indicated in red). The optic chiasm volume (y-axis) for each participant was related to individual variation in measured GC tissue volume (x-axis). Variation in GC tissue volume was modeled as the weighted and combined influence of measured mean GC tissue volume, with and without adjustment for the effects of variation in axial length. The best fit line (with 95% confidence intervals) of the model is shown.

Finally, we examined how well other biometric measures account for optic chiasm size. We find that height (R = 0.36), weight (R = 0.28), axial length (R = 0.27), and age (R = 0.12) all have a weaker relationship with optic chiasm volume than did measures of GCL tissue volume.

## Discussion

In this study we measured normal variation in the profile of the ganglion cell layer of the human retina and related that variation to individual differences in axial length and to the size of the optic chiasm. Our goal is to understand the properties of different measurement units and coordinate frames in characterizing the GCL, and how these choices may be related to counts of the underlying ganglion cell population.

Our study provides a basic empirical result: the population-average profile of the thickness and volume of the GCL along the horizontal meridian. There are limited prior reports of the thickness of the isolated GCL in healthy human eyes, perhaps due to the challenge of segmenting the GC and IP layers in OCT images. Curcio and colleagues (2011) [38] measured the thickness of the GCL along the horizontal meridian in post-mortem histology from the eyes of 18 older donors (ages 40-92 years). Raza and Hood (2015) [15] presented the average, segmented, two-dimensional GCL profile from macular volume scans in 36 healthy participants. Woertz and colleagues (2020) [39] measured the thickness of the segmented GCL along the horizontal and vertical meridians in 25 control participants. All of these measurements identify a peak thickness of approximately 60 microns in the nasal parafoveal region, and a slightly lower peak thickness on the temporal side. Our measurements are in good agreement with this prior work. The current measurements extend further into the periphery of the retina, reaching to 25° eccentricity, as compared to ∼10° eccentricity in prior work. We find that the GC layer thickness plateaus beyond approximately 20°.

It seems reasonable to presume that variation in the size of the ganglion cell layer is related to variation in the total number of retinal ganglion cells. This presumption is well supported for variation across eccentricity. The profile of GCL thickness matches well the profile of retinal ganglion cell counts measured from histological section by Curcio and Allen (1990) [4]. Raza and Hood (2015) [15] used this insight to link their two-dimensional map of GCL thickness to an interpolated map of retinal ganglion cell density. Their method provides an estimate of differences in regional ganglion cell count by examining individual differences in relative GCL thickness.

### Thickness and volume

Variation in eye size, however, complicates efforts to relate GCL thickness to retinal ganglion cell count. A larger eye uses a greater retinal surface area to represent the same size visual field. It may be the case that the same number of retinal ganglion cells are spread over a larger surface area, causing the resulting layer to be thinner. Indeed, we find in our data that the GCL is slightly thinner in larger eyes, consistent with similar, prior measurements of the GC+IPL [3, 32-35].

If the only effect of variation in axial length upon the GCL is to vary the area over which a fixed number of ganglion cells are distributed, then a measurement of tissue volume will correct for this effect. Specifically, there would be no variation by axial length in a measure of cubic mm of GCL tissue per degree square of visual field. We might call this a “volume naïve” version of the stretching model, and we hasten to add that this naivety is used here to motivate a modeling approach for OCT data, but is not to be found in the thoughtful, prior psychophysical work by other investigators. Regardless, our results reject this simple account, as we find instead that larger eyes have a far greater volume of ganglion cell tissue than would be predicted.

One explanation for this effect is that larger eyes have more ganglion cells, and that this greater number of cells results in a greater volume. Psychophysical data argue against this possibility, as angular peripheral spatial acuity is found to not differ across variation in spherical [9, 10]. A related, supportive finding is that, beyond ∼1° eccentricity, the density of cones in eyes of different sizes is invariant with axial length when expressed relative to square degrees of the visual field [40]. Therefore, it seems that variation in spherical refractive error and associated axial length acts to distribute photoreceptors and ganglion cells over different surface areas, but not to alter markedly their numbers.

If the number of cells does not differ in myopic eyes, then either the individual cells are larger, or they are packed less densely, or both. In histologic studies of the central and peripheral nervous system, there is a relationship between number, size, and packing of cells across species, and across differences in body size within species [41]. While these relationships vary substantially by cell type [42], there is a general finding that in larger organs, neural soma size is larger, and the cells are less densely packed.

Overall, our results are consistent with a 10% reduction in ganglion cell density (per mm^3^ of tissue) for each 1 mm increase in axial length.

### Individual differences in cell count and cell size

We modeled the empirical relationship between axial length and GCL tissue volume in our cohort of 50 participants, and then removed this effect from the data. The residuals of this analysis represent individual variation in GCL tissue volume that cannot be explained by variation in axial length. This residual variation could be the result of further individual differences in cell packing—unrelated to axial length—or the consequence of individual differences in the number of retinal ganglion cells in the GCL, or both.

We find that GCL tissue volume, adjusted for axial length, is the best single predictor of individual differences in the size of the optic chiasm in our population. To the extent that GCL and optic chiasm volume reflect the number of retinal ganglion cells and their axons, respectively, this finding identifies axial length-adjusted GCL tissue volume as a measure related to ganglion cell number.

We find an independent and additive prediction of optic chiasm volume by the unadjusted GCL tissue volume. As discussed above, the unadjusted GCL volume measure is dominated by variations in cell size and / or packing that accompany differences in axial length. The correlation with optic chiasm volume suggests that the enlargement of GCL volume in bigger eyes is accompanied by an enlargement of axonal size. This is consistent with myopic eyes having retinal ganglion cells with larger cell bodies, and accompanying larger axons, as has been observed in other neural tissue [43].

While we find GCL tissue volume to be a promising measure to relate to properties of the post-retinal visual pathway, several unanswered questions remain. Even after adjustment for the volumetric effects of axial length, individual differences in adjusted GCL tissue volume combine true variation in cell number with variation in incidental features of cell density. Ideally, we would relate our OCT measures of GCL structure to a more direct measure of retinal ganglion cell count, either through imaging with adaptive optics [44], or through individual difference psychophysical measures that may be related to ganglion cell number [45].

### Distinctions between units for retinal measurement

As has previously been explored for photoreceptor density [40], the current study illustrates the effect of different units of representation for measurements of the GCL, and how these choices interact with variation in the axial length of the eye. Figure 8 presents the modeled GCL profile for three eyes of varying ametropia, in three different measurement frames. We were motivated by theoretical considerations to examine tissue volume per square degree of visual field, expressed as a function of visual field position (Fig 8a). We propose that this coordinate frame is well suited to relate properties of the GCL to visual perception and the post-retinal pathway. The post-retinal visual pathway has a retinotopic organization that is remarkably consistent across individuals when expressed in degrees of polar angle and eccentricity of the visual field [46, 47]. While eyes may vary in shape and size, they have the shared property of representing a fixed amount of the visual field (in degrees) using a collection of retinal ganglion cells of a given volume. A challenge for this coordinate representation is the confounding effect of changes in cell size and packing across axial length. We present here a model of this effect, which allows us to remove its influence.

**Figure 8.**
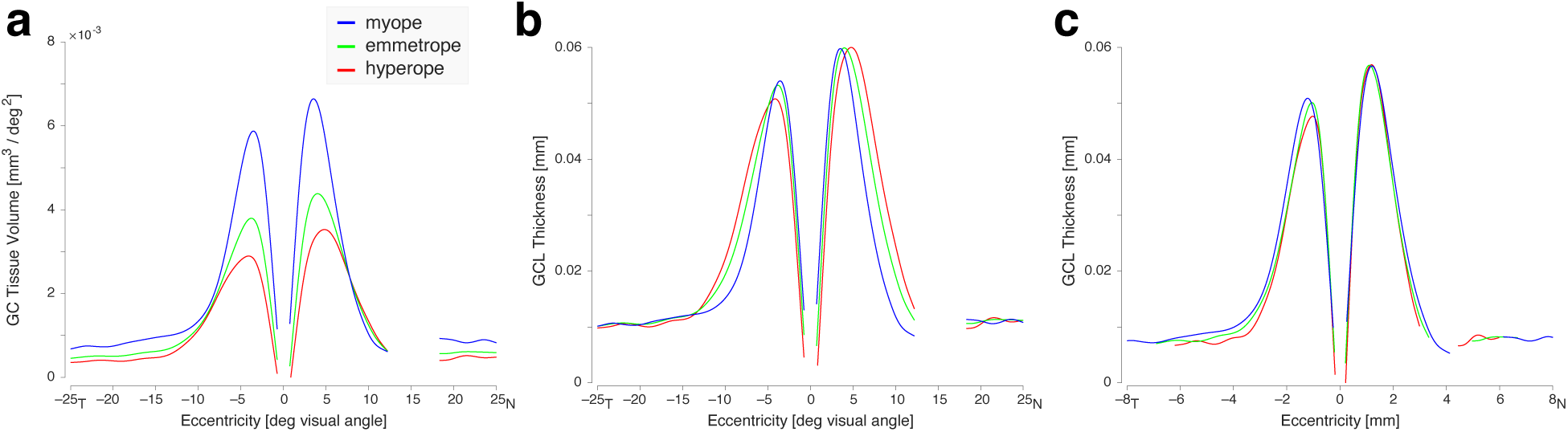
The retinal ganglion cell layer considered in different coordinate frames. The profile of the GCL along the horizontal meridian is modeled for eyes of varying axial lengths (myope 27.57 mm; emmetrope 23.58 mm; hyperope 21.79 mm). This same set of profiles is plotted for three different choices of unit and coordinate frames. (a) GC tissue volume per square degree of visual angle as a function of eccentricity in degrees of visual angle; (b) GCL thickness in mm as a function of eccentricity in degrees of visual angle; (c) GCL thickness in mm as a function of eccentricity in mm of retina. For the last of these, the extent of the plotted functions along the x-axis varies with eye size.

Measurements of the GCL and related structures have traditionally been made as thickness (in mm or microns), expressed as a function of retinal distance in degrees of visual angle (Fig 8b) or mm (Fig 8c). The choice of x-axis units obviously interacts with spherical ametropia, as the size of the vitreous chamber influences the conversion of degrees to mm. Prior work has carefully characterized normal variation in the shape of the foveal pit, and how that variation is related to axial length [48-50]. As the morphology of the foveal pit is influenced by multiple retinal layers, the GCL thickness profiles we report here cannot be directly related to these prior studies. Indeed, we note that Woertz and colleagues (2020) [39] found that the GC and IP layers had quite variable relative thicknesses across individuals, leading us to be cautious in attempting to relate our measurements of a single retinal layer to larger properties of retinal morphology.

### Conclusion

Several neurologic [51, 52] and ophthalmologic [53, 54] diseases are associated with the loss of retinal ganglion cells. In these conditions, OCT measures of retinal structures have been pursued as imaging biomarkers of the health of retinal cells [55-58]. Existing analyses, however, rarely account for the effect that variation in eye size has upon OCT measures. In the current study we characterize the relationship between variation in axial length and measurements of the ganglion cell layer. Based upon an assumption that eyes of different sizes do not systematically vary in their total number of retinal ganglion cells, we introduce a measure of the volume of ganglion cell tissue relative to the visual field, and an adjustment for axial length to remove variation in cell size and packing. The resulting measure accounts for individual differences in the size of the optic chiasm, supporting its use to characterize the post-retinal visual pathway.

## Acknowledgments

Supported by National Institutes of Health Grant U01EY025864: Human Connectomes in Low Vision, and P30 EY001583: Core Grant for Vision Research.

## Competing interests

The authors have no competing interests to declare.

## Notes

https://github.com/gkaguirrelab/retinaTOMEAnalysis

### Competing Interest Statement

The authors have declared no competing interest.

